# Neurologically healthy humans’ ability to make saccades toward unseen targets

**DOI:** 10.1101/2022.05.27.493699

**Authors:** Henri Olkoniemi, Mikko Hurme, Henry Railo

## Abstract

Some patients with a visual field loss due to a lesion in the primary visual cortex (V1) can shift their gaze to stimuli presented in their blind visual field. The extent to which a similar “blindsight” capacity is present in neurologically healthy individuals remains unknown. Using retinotopically navigated transcranial magnetic stimulation (TMS) of V1 (Experiment 1) and metacontrast masking (Experiment 2) to suppress conscious vision, we examined neurologically healthy humans’ ability to make saccadic eye movements toward visual targets that they reported not seeing. In the TMS experiment, the participants were more likely to initiate a saccade when a stimulus was presented, and they reported not seeing it, than in trials which no stimulus was presented. However, this happened only in a very small proportion (~8%) of unseen trials, suggesting that saccadic reactions were largely based on conscious perception. In both experiments, saccade landing location was influenced by unconscious information: When the participants denied seeing the target but made a saccade, the saccade was made toward the correct location (TMS: 68%, metacontrast: 63%) more often than predicted by chance. Signal detection theoretic measures suggested that in the TMS experiment, saccades toward unseen targets may have been based on weak conscious experiences. In both experiments, reduced visibility of the target stimulus was associated with slower and less precise gaze shifts. These results suggest that saccades made by neurologically healthy humans may be influenced by unconscious information, although the initiation of saccades is largely based on conscious vision.

## INTRODUCTION

Humans orient themselves toward stimuli using fast eye movements called saccades (e.g., Rayner, 1998). This process is automatic, and it is often assumed that it can be initiated in response to stimuli that are not consciously seen (e.g., McCormick, 1997). Some patients with lesions in the primary visual cortex (V1) have been shown to retain their ability to make saccades toward stimuli that they report not seeing (Cowey, 2010; Pöppel et al., 1973; Weiskrantz et al., 1974). However, the extent to which this ability reflects completely unconscious perception and is present in neurologically healthy individuals remains open (see Railo and Hurme, 2021 for a review). In the present study, we used transcranial magnetic stimulation (TMS) of V1 (Experiment 1) and visual metacontrast masking (Experiment 2) to test whether neurologically healthy participants could initiate saccadic eye movements toward stimuli they reported not seeing.

According to a popular model, visual pathway involving the V1 mediates conscious perception of visual stimuli. Phylogenetically older visual pathways via the pulvinar and superior colliculus (SC) are often assumed sufficient for triggering behavioral responses toward visual stimuli that are not consciously registered (Danckert and Rossetti, 2005; Schneider, 1969; Tamietto and De Gelder, 2010; Baldwin et al., 2017). Given that saccadic eye movements are initiated by neurons in the SC (Wurtz, 2010), saccades are prime candidates for actions that can be triggered by unconscious stimuli. In nonhuman primates, V1 lesions lead to spatially more imprecise saccades (Yoshida et al., 2008). Yet, despite the lesion, saccadic reaction times to stimuli in the affected hemifield are relatively fast and show less variance when compared to saccades in the intact visual field (Yoshida et al., 2008), suggesting that they are reflexive behaviors. Studying the extent to which saccadic eye movements are associated with conscious experience of the stimuli is particularly difficult in nonhuman primates, but evidence suggests that despite V1 lesions, monkeys may retain a severely degraded form of conscious visual perception of stimuli (Isa and Yoshida, 2021; Yoshida and Isa, 2015).

In humans, V1 lesions typically cause blindness in the corresponding visual field but stimuli that are presented in the affected visual field may still influence the behavior of these patients, which is known as *blindsight* (Cowey, 2010; Pöppel et al., 1973; Weiskrantz et al., 1974). It is debated whether blindsight is based on completely unconscious visual information (Michel and Lau, 2021; Weiskrantz, 2009), or whether it is a form of severely degraded conscious vision (Campion et al., 1984; Ffychte and Zeki, 2011; Phillips, 2021).

Oculomotor responses to stimuli in the blind visual field have been studied relatively extensively in blindsight patients. Pöppel et al. (1973) observed that although patients (*N* = 4) with V1 lesions could not accurately shift their gaze to stimuli presented in the blind field, the endpoints of the saccades were biased by the stimulus presented to the blind visual field across trials. Weiskrantz et al. (1974) observed that a blindsight patient could accurately fixate targets presented 5° from fixation in the blind visual field, but the accuracy of saccades to more eccentric targets was at chance level. Perenin and Jeannerod (1978) examined six hemidecorticated patients and observed that “the eyes were turned toward the target each time it had to be located” (p. 8), although this could only be precisely measured in two patients. Similarly, Zihl (1980) studied three patients and observed that two of them could shift their gaze to stimuli presented in the blind visual field. Interestingly, in Zihl’s (1980) study, saccades to blind visual field locations were initiated about 100 ms faster than saccades to targets presented in the intact visual field, suggesting that they were reflexive and not based on a conscious decision. Barbur et al. (1988) observed that a patient with a left visual cortical lesion could initiate rapid saccades toward visual stimuli in the affected hemifield. Rafal et al. (1990) showed that saccades toward the intact visual field could be slowed down by stimuli presented in the blind visual field. This slowing down was caused only by distractors presented in the (monocular) temporal visual field, suggesting that the effect was mediated by the visual pathway to SC. Savina and Guitton (2018) studied two hemidecorticated patients and reported that patients’ saccades toward the blind visual field were biased by stimuli presented in the blind field.

The above findings suggest that some patients with blindsight can fixate targets presented in their blind visual field, but there is variation between patients. The individual differences could be due to variation in the extent of the lesions. Further, blindsight may be partly a learned ability. For example, Zihl (1980) and Roth et al. (2009) observed that training improved patients’ ability to fixate targets in the blind visual field. Blindsight could also be due to the neural reorganization of the white matter pathways after the lesion. Nonhuman primates (Isa and Yoshida, 2021) and some patients with blindsight (Bridge et al., 2008; Leh, Johansen-Berg, et al., 2006; Leh, Mullen, et al., 2006; Tamietto et al., 2012) show anatomical connections that are not present in patients without blindsight or in neurologically healthy controls. This means that studies on blindsight patients leave open the question of the extent to which V1 contributes to gaze shifts in neurologically healthy individuals (Railo and Hurme, 2021).

Early visual cortex (V1–V3) TMS suppresses the visibility of a briefly presented stimulus (Amassian et al., 1989; de Graaf et al., 2014). This allows testing of whether blindsight-like behavior can be detected in neurologically healthy individuals (known as “TMS-induced blindsight”). Although studies do not provide support for a completely unconscious TMS-induced blindsight of visual features such as color or orientation (Hurme et al., 2020; Koivisto et al., 2021; Lloyd et al., 2013; Railo et al., 2014), neurologically healthy participants may be able to detect the presence or roughly localize a visual stimulus they report not consciously seeing (Christensen et al., 2008; Hurme et al., 2017; Railo and Koivisto, 2012; Ro, 2008, reviewed in, Railo and Hurme, 2021). Ro et al. (2004) reported that distractor stimuli whose visibility was suppressed due to TMS delayed saccadic eye movements to visible targets. However, a test of neurologically healthy individuals’ ability to initiate a saccade toward a target whose conscious visibility has been suppressed by V1 TMS is lacking.

The aim of the present study was to determine whether saccadic eye movements can be initiated toward targets that participants report not consciously seeing. We presented visual stimuli to the left or right visual field and asked the participants to rapidly shift their gaze to the target. The participants then rated how well they saw the stimulus. In Experiment 1, visibility of the stimulus was suppressed by TMS to the left or right V1. If the saccade control system can initiate gaze shifts without V1, the participants should be able to shift their gaze toward the target location even when stimulus visibility is suppressed by V1 TMS. In Experiment 2, stimulus visibility was suppressed using visual metacontrast masking. Metacontrast masking is assumed to inhibit target-related feedback activity but leave feedforward activity (in V1, and other areas) largely intact (Breitmeyer and Ogmen, 2010; Macknik and Livingstone, 1998; Railo and Koivisto, 2012). Consequently, in Experiment 2, we expect that the participants should be able to make saccades toward the reportedly unseen targets if, in general, it is possible to make saccades toward unseen targets.

## EXPERIMENTAL PROCEDURES

### Participants

Twelve University of Turku students between the ages of 21–46 (11 women, *M*_age_ = 27, *SD*_age_ = 7) participated in Experiment 1. The visual areas of these individuals were mapped using functional magnetic resonance imaging (fMRI). All participants had normal or corrected-to-normal vision. Participants were paid a small monetary fee for taking part in the experiment.

Thirteen University of Turku students (12 women, *M_Age_* = 23, *SD_Age_* = 3.70) took part in Experiment 2 for course credit. The participants in Experiment 2 did not take part in the first experiment. All participants had normal or corrected-to-normal vision.

The study was approved by the ethics committee of the Hospital District of Southwest Finland and was conducted in accordance with the Declaration of Helsinki and with the understanding and written consent of each participant.

### Apparatus

#### Magnetic Resonance Imaging

For the participants who took part in Experiment 1, MRI was performed in Turku PET Center using 3T Philips MRI. A high-resolution (voxel size = 1 mm^3^) T1-weighted anatomical image of the whole head was acquired (3D TFE) for each participant. Visual cortical areas were mapped with fMRI using a modified version of the multifocal procedure (Henriksson et al., 2012) described by Vanni et al. (2005). The multifocal visual stimuli were presented in the scanner using VisualSystem HD (NordicNeuroLab, Bergen, Norway) video goggles (1920 × 1200 resolution) and Presentation software (Neurobehavioral Systems, Inc., Albany, CA, USA). For the retinotopic mapping, the major imaging parameters were repetition time of 1,800 ms, echo time of 30 ms, flip angle of 60°, field of view of 250 mm, matrix of 96 × 96, and a 2.5 mm^3^ voxel size. Twenty-nine slices were acquired in interleaved order. Standard preprocessing with slice time and motion correction was followed by estimation of the general linear model with the SPM8 Matlab toolbox.

#### Transcranial Magnetic Stimulation

In Experiment 1, TMS was delivered using a MagVenture MagPro X100 stimulator and a liquid-cooled 65 mm figure-of-eight coil (Cool-B65). TMS intensity was set to 85% of the maximum stimulator output. Stimulation location was neuronavigated based on the individual participants’ MRIs using the Localite TMS Navigator 3.0.48 navigation system. The targeted stimulation areas in V1 were defined individually based on retinotopic maps of the visual cortex. The two stimulation targets were the left and right hemisphere V1 locations, corresponding to the receptive fields that process the location where the visual stimulus was presented. Anatomically, these correspond to locations in the upper bank of the calcarine sulci. These V1 locations were selected because they are easiest to stimulate using TMS. The left and right hemisphere locations were stimulated in alternating experimental blocks (order counterbalanced).

#### Eye Tracking

In both Experiments 1 and 2, eye movements were recorded monocularly using EyeLink 1000 Plus (SR Research Ltd., Ontario, Canada) at a sampling frequency of 1000 Hz. The participants were seated 82 cm from the screen, and a chinrest was used to stabilize the head. The eye tracker was set up and calibrated using a three-point calibration screen (successful calibration was determined by *M_error_* < 0.5° in visual angle, error at each point < 1°).

### Stimuli and Tasks

#### Experiment 1

In Experiment 1, participants were presented with visual targets 2° from the fixation cross (which was in the center of the monitor) in the bottom left and right quadrants in random order. The target stimuli were dots presented against a light gray background (75 cd/m^2^) for one screen refresh (8.3 ms). The stimuli size was set to 0.20° but increased if the participant reported that they saw the target in less than 80% of the practice blocks ran before the actual experiment (stimulus size *M* = 0.22°, *SD* = 0.02°). The luminance of the target stimulus was 51 cd/m^2^. These visual stimulus parameters roughly correspond to those used by Railo and Koivisto (2012). The experiment was run in MATLAB (version R2014b), and the target stimuli were presented on a 24” VIEWPixx /Lite LCD-monitor with refresh rate of 120 Hz and a resolution of 1920 × 1080 pixels.

The TMS pulse was delivered 100 ms after the onset of the visual target stimuli in each trial, as previous studies have shown that this stimulus onset asynchrony (SOA) reliably suppresses conscious perception (de Graaf et al., 2014). Moreover, Hurme et al. (2017) suggested that earlier TMS SOAs may disturb unconscious visual processing by interfering with feedforward visual activity in V1. Given that the target was presented to either the left or the right visual field, the stimulated cortical area was either contralateral or ipsilateral with respect to the visual target stimulus. The contralateral trials were experimental trials in which strong suppression by TMS was expected, and the ipsilateral trials were control trials.

The participants indicated where the target was presented by moving their eyes as fast as possible to the location where the target appeared or keeping their eyes in the central location if they did not perceive any target. Participants were given 900 ms to give their answers. Then, the participants rated the subjective visibility of the target stimuli on a four-step scale (Ramsøy and Overgaard, 2004; Sandberg et al., 2010). The definitions of the four alternatives were carefully explained to the participants. The lowest alternative (0) indicates that the participants *did not see the target at all*. The second lowest option (1) indicates that the participants felt that they *detected a small glimpse of the target*. During the instructions, it was emphasized that if the participants felt that they saw anything related to the target stimulus (no matter how weak), they should always choose the second lowest option. That is, the lowest alternative was reserved only for trials in which the participants felt confident that they did not see the target at all (throughout the manuscript, we refer to these trials as “unseen” or “unconscious”). The third option (2) indicates that the participant *perceived the target pretty well, but not perfectly*, and the highest rating (3) corresponded to the *perfectly seen target*. The responses were given using a numeric keypad on a keyboard.

Before the actual experiment, each participant completed a practice session to familiarize them with the task and determine whether the stimulus size was adequate for them. The practice session contained blocks of 20 trials. The practice blocks were repeated until the participants felt comfortable with the task and could correctly indicate the target location in more than 80% of the trials. On average, the participants completed two practice blocks before the actual experiment. In the experiment, the target stimuli were presented in blocks of 70 trials. We aimed to complete at least eight experimental blocks (i.e., 560 trials) with each participant within a 120-minute experimental session. On average, the participants completed 8.25 blocks, or 577.50 trials (*SD_blocks_* = 0.87, *SD*_trials_ = 60.62). Each experimental block included 20% of the catch trials (i.e., no target stimulus was presented). The experimental setup and the simulated area are presented in Figure 1A and 1B.

**Figure 1.**
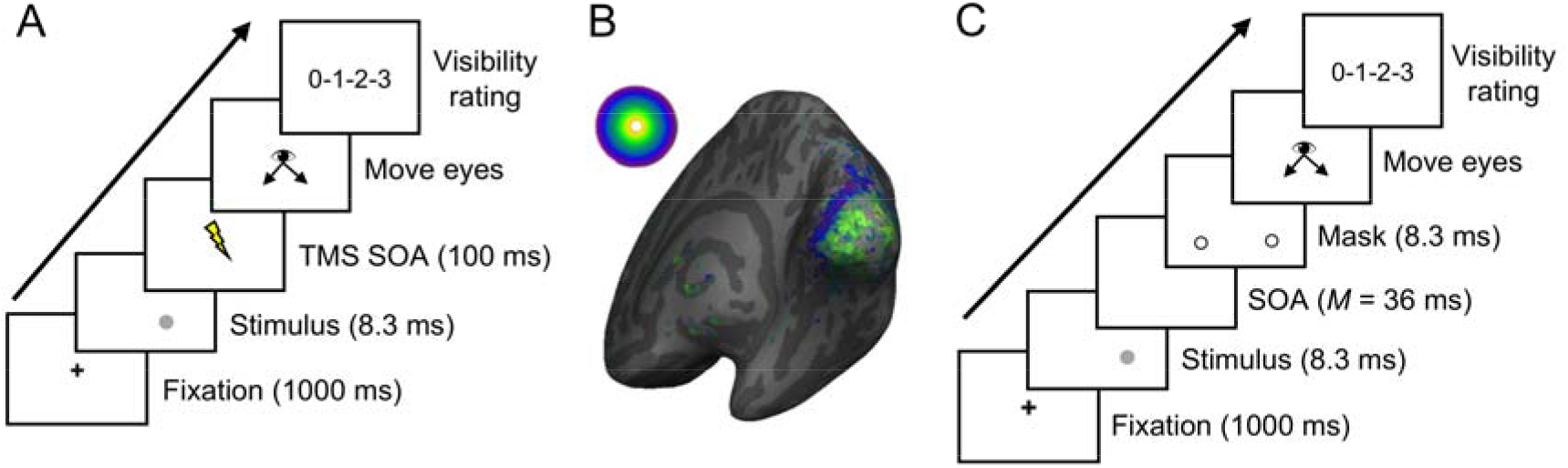
Schematic presentation of the target stimuli and the experimental procedures. (A) Timeline of the experimental trial in Experiment 1. The participants’ tasks were to move their eyes as fast as possible to the location where the target appeared or keep their eyes in the central location if they did not perceive any target, and then to report their degree of confidence in this decision. The visibility of the target stimulus was manipulated using transcranial magnetic stimulation (TMS). (B) An example of retinotopically mapped visual cortical areas used to determine the stimulated area in Experiment 1. Here, color represents eccentricity, overlayed on the inflated right hemisphere of one participant (visualized using FreeSurfer software). (C) Timeline and the experimental procedure used in Experiment 2. The experimental procedure was similar to Experiment 1 except the visibility of the stimulus was manipulated using metacontrast masking.

#### Experiment 2

Experiment 2 differed from Experiment 1 only in the use of metacontrast masking to manipulate the visibility of the target stimulus. The mask comprised of two black annuli (line width = 0.07°). The mask had the same inner diameter as the outer diameter of the target. The size of the target stimuli was set to 0.20°, as this stimulus size was perceived well by the participants during the practice trials in Experiment 1. The mask was presented for one screen refresh, and the SOA between the target and mask onset was determined during the practice trials. The SOA was set so that each participant would be able to consciously perceive about the same number of targets as in Experiment 1. The SOA was, on average, 36 ms *(SD =* 9 ms). During the experimental session, on average, 7.9 blocks were presented to the participants, with each block comprising 50 trials (the total number of trials was, on average, 396.15 trials per participant, *SD_blocks_* = 1.12, *SD_trials_* = 55.76). The schematic presentation of the stimuli and the experimental procedure used in Experiment 2 are illustrated in Figure 1C.

### Procedure

Upon arrival, the participants were naïve to the purpose of the experiment, and the specific nature of the task was explained to them only after the experiment. First, each participant signed an informed consent form. After that, the eye-tracking and, in Experiment 1, TMS systems were introduced to them. Next, the experimental procedure was explained. The participants then completed practice blocks, and after these blocks, the actual experiment began. The experimental session lasted about 120 min. There was always a mandatory break in the middle of the experiment to maintain the participants’ vigilance, and they were offered refreshments. In addition, the participants were given the opportunity to rest after each block.

### Data Preparation and Analysis

Saccadic answers were determined for each trial as follows. Eye movement was determined as a saccade using 30°/s threshold for velocity and 8000°/s^2^ for acceleration (Stampe, 1993). For the saccade to be determined as an answer, it needed to cross the vertical and horizontal halfway point between the fixation cross and stimulus location (i.e., >1° from the fixation cross). These thresholds are illustrated in Figures 2A and 6A with dashed lines. If multiple saccades were present during the trial, only the first saccade after stimulus presentation that crossed these thresholds was considered. This procedure yields a binary measure of whether the participant made a saccade to *either* target location. We call this measure a “saccadic reaction.” This enables us to examine whether visual stimuli participants report not seeing trigger saccades, irrespective of whether the gaze shift was toward the correct location. In 19% of the trials in Experiment 1 and 7% in Experiment 2, the saccade was made before stimulus onset; consequently, these trials were removed from the data (the frequency and proportion of saccadic reactions for Experiment 1 are presented in Table 1, and in Table 5 for Experiment 2).

**Figure 2.**
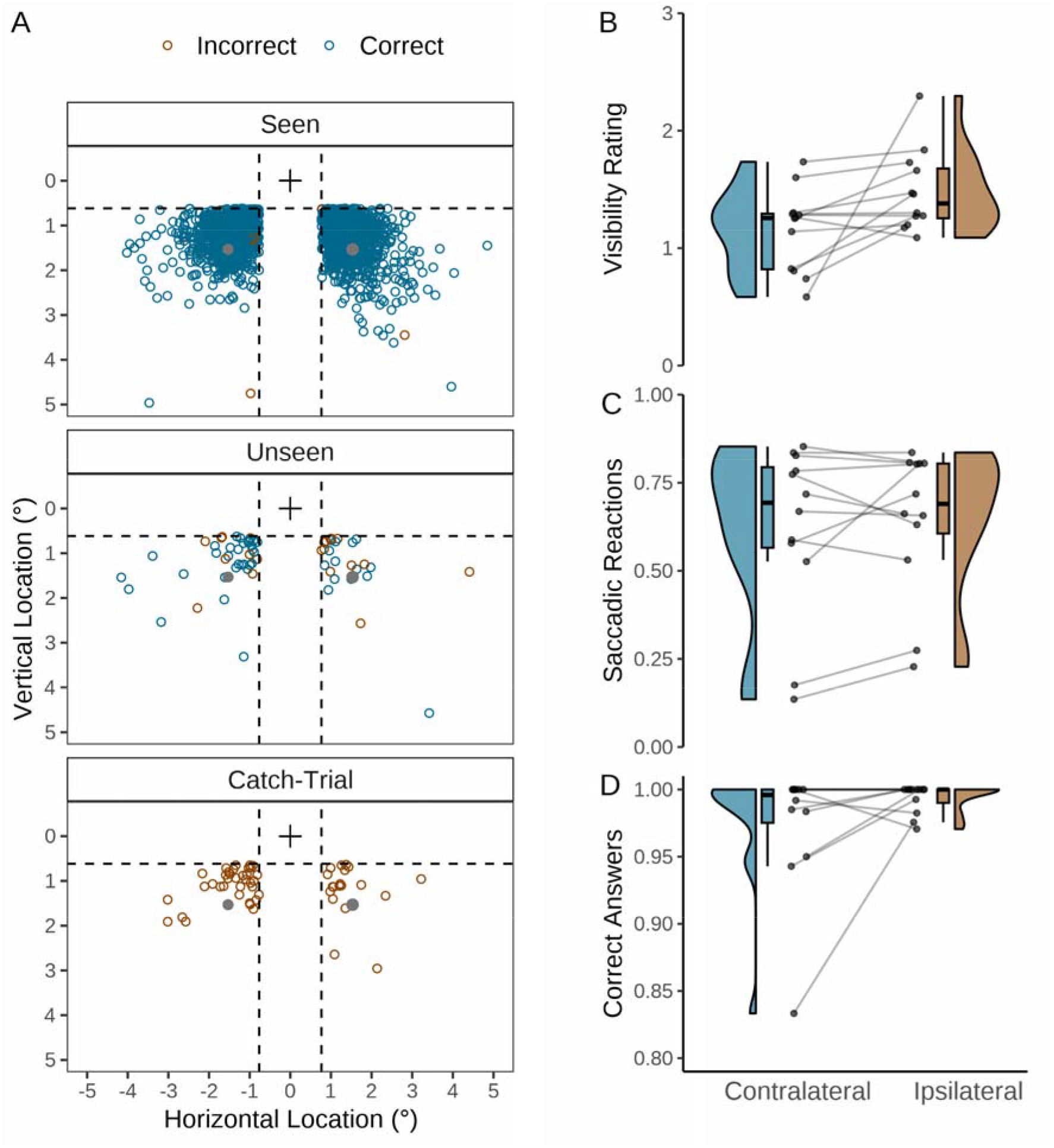
Saccadic eye movements in different experimental conditions. (A) All landing locations of the saccadic reactions. The X-axis and Y-axis present landing position in the visual angle (°). The black cross indicates the fixation point location, and the gray dots indicate the target stimulus locations. Dashed lines indicate horizontal and vertical thresholds used to determine whether the saccade was counted as a reaction or not. (B) Average visibility ratings in contralateral and ipsilateral conditions. (C) Proportion of saccadic reactions in contralateral and ipsilateral conditions. (D) Proportion of correct saccadic answers (of the saccadic reactions made) in contralateral and ipsilateral conditions. In panels B–D, the lines in the middle represent the direction of change for each participant between the conditions.

**Figure 3.**
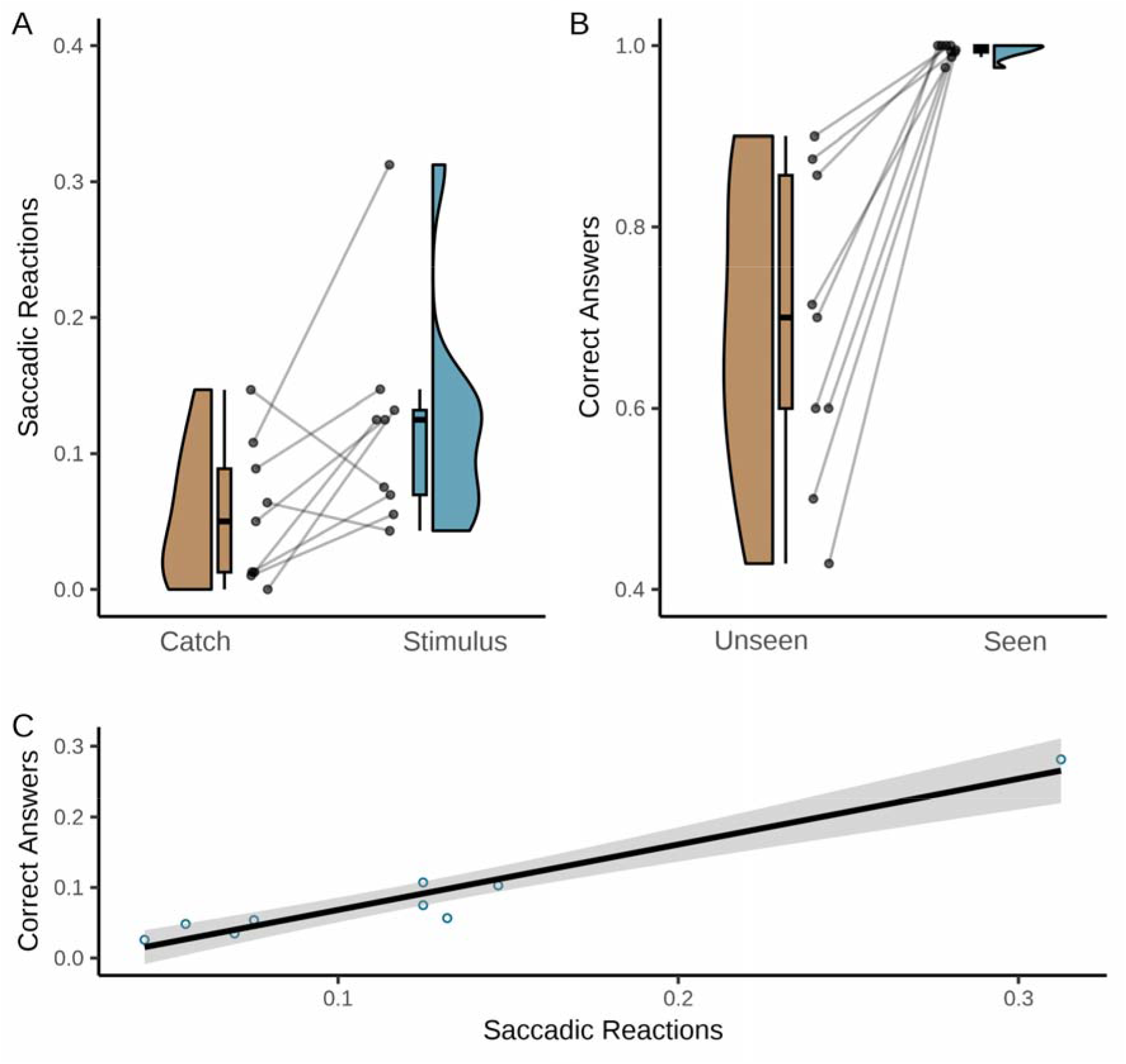
Saccadic reactions as a function of conscious perception of the target. The lines represent the direction of the effect for each participant. (A) Proportion of saccadic reactions (to either correct or incorrect target location) when the participants reported not seeing the target at all (i.e., visibility rating = 0). (B) Proportion of correct saccadic answers (in relation to total number of saccadic reactions) when the participants reported seeing (visibility rating > 0) or not seeing the target (visibility rating = 0). In panels B and C, the lines in the middle represent the direction of change for each participant between the conditions. (C) Relationship between the proportion of saccadic reactions to unseen trials and the proportion of correct saccadic answers (i.e., toward the target location; in relation to number of unseen trials). Shaded area represents 95% CI.

**Figure 4.**
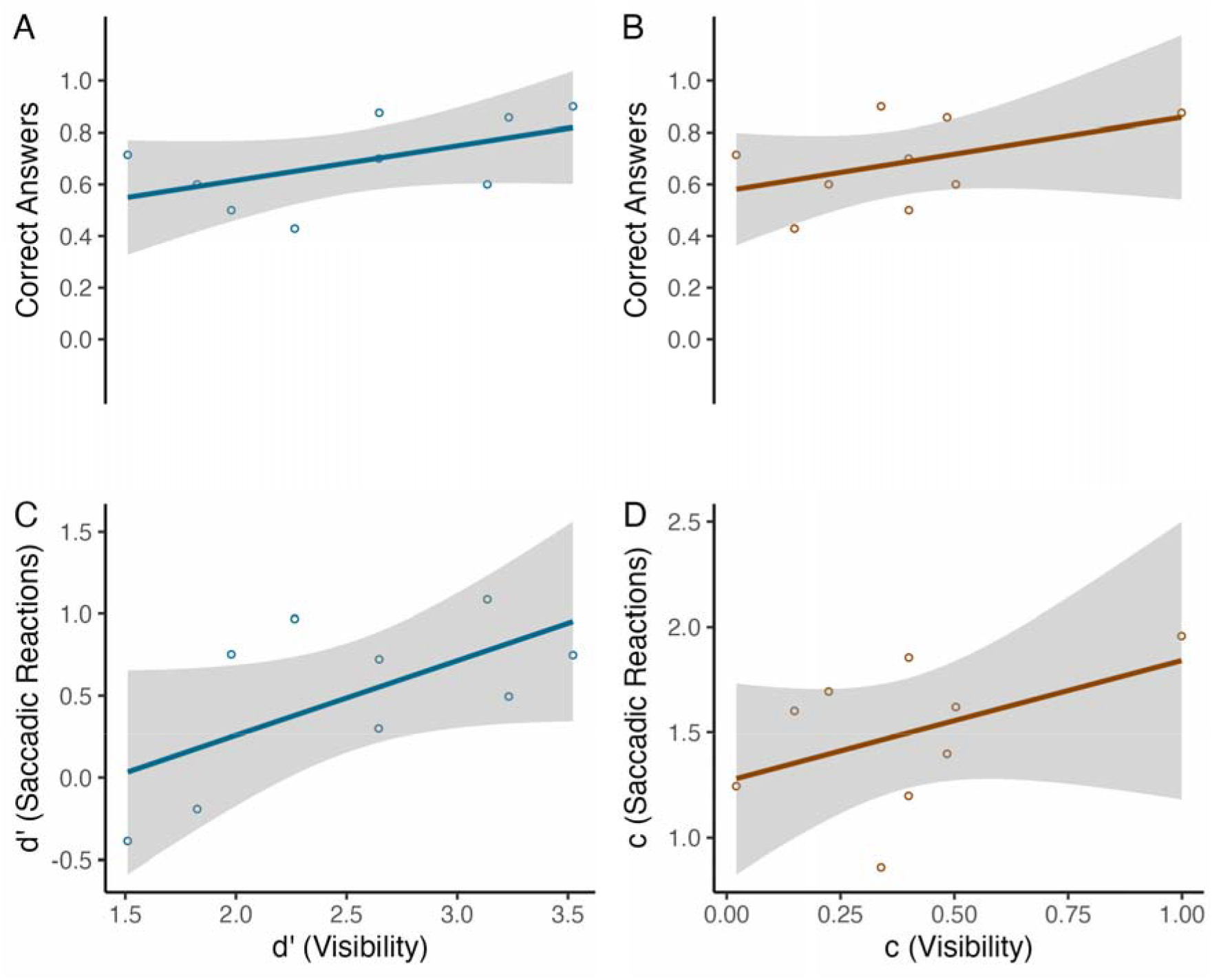
Association between saccades made in response to unconscious targets, and sensitivity to consciously detect the stimulus. (A) The correlation between the sensitivity of consciously detecting the stimulus, and the proportion of correct saccadic answers (of all the saccadic reactions) in unconscious trials. (B) The correlation between the criterion for reporting conscious perception, and the proportion of correct saccadic reactions (of all the saccadic reactions) in unconscious trials. (C) The correlation between the sensitivity of subjectively detecting the stimulus, and the sensitivity of reacting to the stimulus. (D) The correlation between the criterion for reporting conscious perception, and the criterion for making a saccadic reaction. In all panels, the gray area represents the 95% CI.

**Figure 5.**
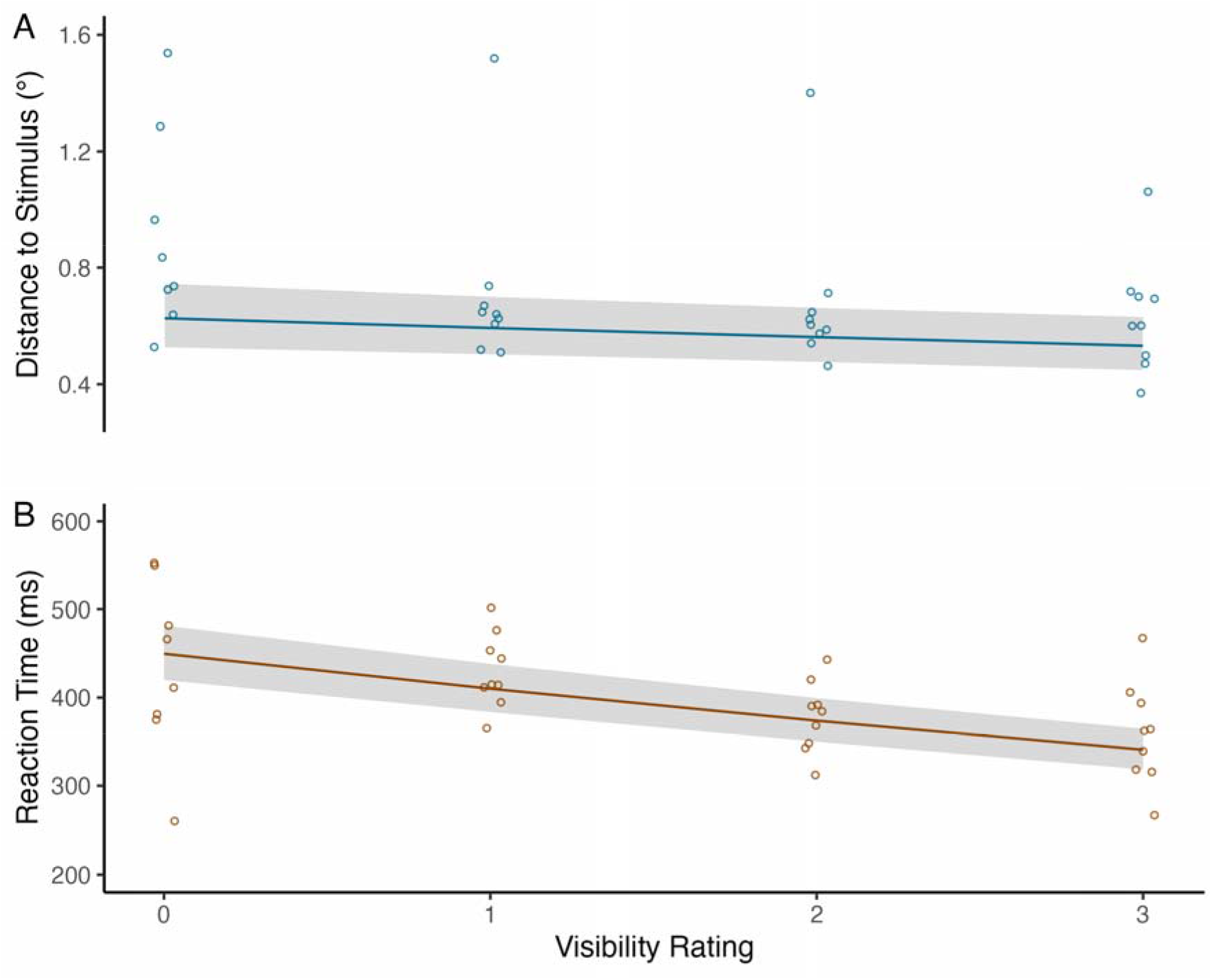
Precision of saccadic reactions and saccadic reaction times. (A) Model estimates for the effect of visibility on saccade accuracy. (B) Model estimates for saccadic reaction times. For the purpose of illustration, the model estimates were back-transformed from the log values in both panels. The shaded area represents the 95% CI. Circles represent the observed means for each participant.

**Figure 6.**
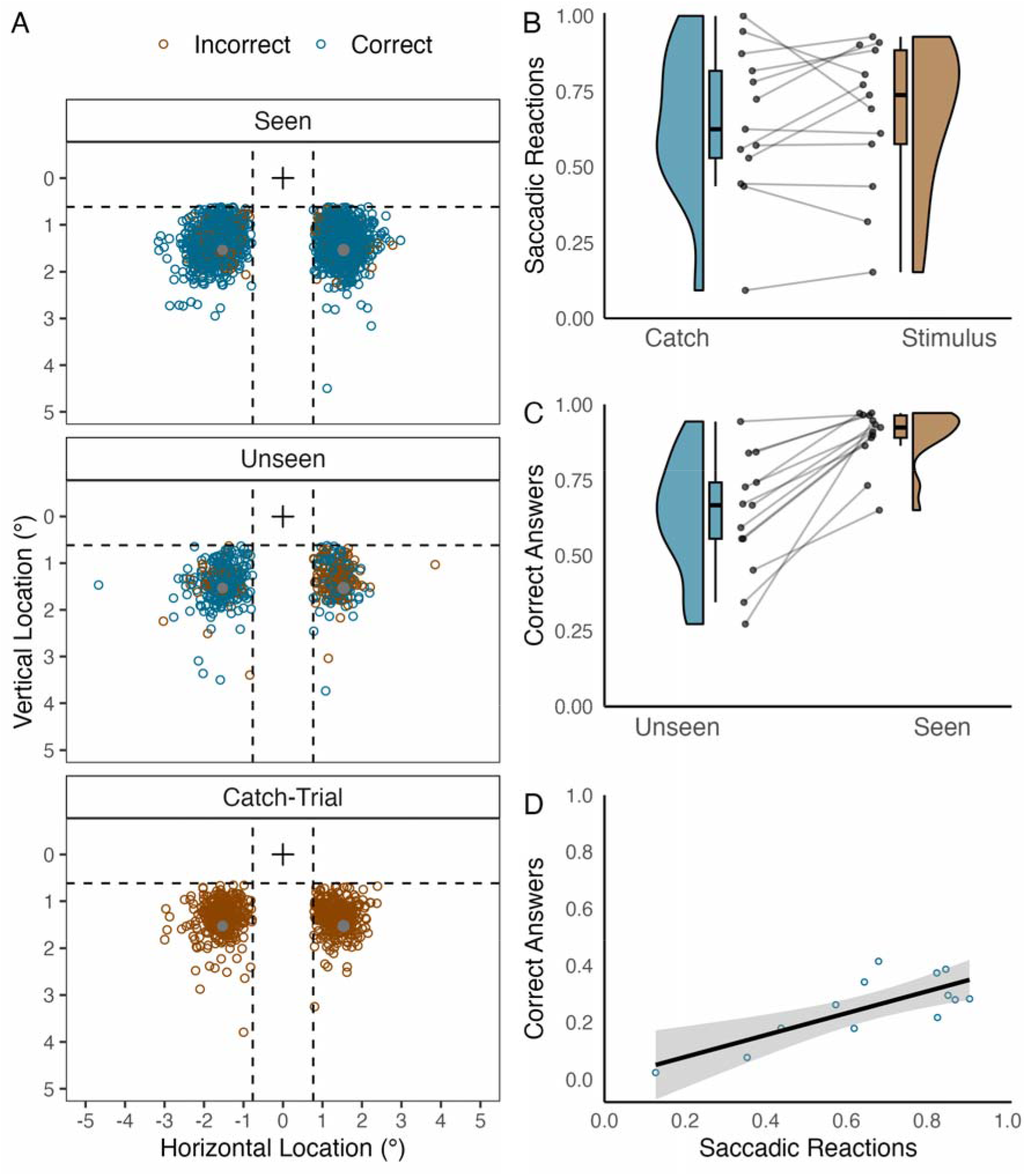
Saccades to targets in different experimental conditions. (A) All the landing locations of the saccadic reactions. The X-axis and Y-axis present landing position in the visual angle (°). The black cross indicates the fixation point location, and the gray dots indicate the target locations. Dashed lines indicate the horizontal and vertical thresholds used to determine whether the saccade was counted as an answer. (B) Proportion of saccadic reactions (to either stimulus location) when participants report not seeing the target at all (i.e., visibility rating = 0). (C) Proportion of correct saccadic answers (in relation to total number of saccadic reactions) when participants reported seeing (i.e., visibility rating > 0) and not seeing the target (i.e., visibility rating = 0). (D) Relationship between the proportion of saccadic reactions to unseen trials, and the proportion of correct saccadic answers (i.e., toward the target location; in relation to number of unseen trials). Shaded area represents 95% CI.

**Table 1:** Frequency and the proportion of saccadic reactions across experimental conditions

| Visibility | Trial | TMS | N<br>(total) | Saccadic<br>Reactions |  |  | Correct Answers |  |  |
| --- | --- | --- | --- | --- | --- | --- | --- | --- | --- |
|  |  |  |  | N | M | SD | N | M | SD |
| Unseen | Catch |  | 765 | 33 | .04 | .20 | - | - | - |
|  | Stimulus | Contralateral | 534 | 38 | .07 | .27 | 21 | .53 | .24 |
|  |  | Ipsilateral | 359 | 31 | .09 | .28 | 26 | .85 | .21 |
| Seen | Catch |  | 81 | 21 | .26 | .44 | - | - | - |
|  | Stimulus | Contralateral | 1674 | 1105 | .66 | .47 | 1094 | .97 | .05 |
|  |  | Ipsilateral | 2172 | 1454 | .67 | .47 | 1449 | .99 | .01 |

**Table 2.** Descriptive statistics for precision and reaction times of the correct saccadic reactions

| TMS | Visibility | Distance from the stimulus location |  | RT (ms) |  |
| --- | --- | --- | --- | --- | --- |
|  |  | *Mean* | *SD* | *Mean* | *SD* |
| Contralateral | Unseen | 1.20 | 1.07 | 426 | 148 |
|  | Seen | 0.65 | 0.38 | 389 | 93 |
| Ipsilateral | Unseen | 0.82 | 0.66 | 431 | 133 |
|  | Seen | 0.64 | 0.61 | 395 | 90 |

*Note.* The distance from stimulus location is reported as visual degrees.

**Table 3.** Model for the precision of the saccadic reaction

| **Random Effects** |  | *n* | *Variance* | *SD* |
| --- | --- | --- | --- | --- |
| Participant (intercept) |  | 9 | 0.06 | 0.25 |
| Participant (TMS location) |  |  | 0.0003 | 0.02 |
| Residual |  |  | 0.34 | 0.58 |
| **Fixed Effects** |  | *Estimates* | *95% CI* | *t* |
| Intercept |  | -0.57 | -0.74 – -0.41 | **-6.77** |
| TMS Location (contralateral vs. ipsilateral) |  | -0.02 | -0.04 – 0.03 | -0.70 |
| Visibility |  | -0.04 | -0.06 – -0.01 | **-2.73** |
| TMS Location (contralateral vs. ipsilateral) × Visibility |  | 0.01 | -0.04 – 0.06 | 0.40 |

*Note.* $t > |1.96|$ are bolded. $N$ of observations = 2097.

**Table 4.** Model for saccadic reaction times.

| Random Effects | *n* | Variance | SD |
| --- | --- | --- | --- |
| Participant (Intercept) | 9 | 0.01 | 0.10 |
| Participant (TMS location) |  | 0.002 | 0.05 |
| Residual |  | 0.04 | 0.19 |
| Fixed Effects | Estimates | 95% CI | *t* |
| Intercept | 5.96 | 5.89 – 6.03 | **176.10** |
| TMS Location (contralateral vs. ipsilateral) | 0.01 | -0.02 – 0.05 | 0.65 |
| Visibility | -0.07 | -0.08 – -0.06 | **-15.78** |
| TMS Location (contralateral vs. ipsilateral) ×<br>Visibility | -0.00001 | -0.02 – 0.02 | -0.01 |

*Note.* $t > |1.96|$ are bolded. *N* of observations = 2037

**Table 5.** Frequency and proportion of saccadic reactions across experimental conditions

| Visibility | Trial | *N*<br>(total) | Saccadic Reactions |  |  | Correct Answers |  |  |
| --- | --- | --- | --- | --- | --- | --- | --- | --- |
|  |  |  | *N* | *M* | *SD* | *N* | *M* | *SD* |
| Unseen | Catch | 555 | 337 | .61 | .49 | - | - | - |
|  | Stimulus | 793 | 520 | .66 | .48 | 354 | .63 | .20 |
| Seen | Catch | 353 | 280 | .79 | .41 | - | - | - |
|  | Stimulus | 3087 | 2639 | .85 | .35 | 2375 | .89 | .10 |

Another binary measure was created to indicate whether the saccadic reaction was done in the direction of the stimulus (i.e., correct answer) or in the opposite location (i.e., wrong answer). This measure was used to calculate the mean proportion of correct saccadic answers for each participant. We also calculated the distance in visual degrees between the saccade end location and the target location (called “saccadic precision”). This gives us a continuous measure of saccade accuracy. Lastly, saccadic reaction times were calculated relative to the onset of the stimulus.

Data were analyzed with R statistical software (Version 4.1.2; R Core Team, 2021). The main analyses focused on trials in which participants reported not seeing the target at all. We assessed whether participants were able to make saccadic reactions more often when a stimulus was present than when it was not. We also determined whether the participants were able to shift their gaze toward the correct target location more often than predicted by chance (one-sample t-test against 50%). We assessed *saccadic reactions to unconscious targets* by calculating signal detection theoretic sensitivity (d’) and response criterion (c) based on the z-scores of hit and false alarm rates obtained through the inverse cumulative distribution function (Macmillan and Creelman, 2005). A hit was defined as a saccade landing to either stimulus location when a stimulus was presented (but reported not seen). False alarms were trials in which the participant made a saccade to either target location when no stimulus was presented (and visibility rating = 0). The participants’ sensitivity (d’) and criterion (c) for *reporting a conscious perception* of the target were calculated. A hit was defined as a visibility rating > 0 when a stimulus was presented. False alarms were trials in which the participant reported seeing a target (rating > 0) when no stimulus was presented.

*Saccadic precision* and *saccadic reaction times* were analyzed with linear mixed-effects models using the *lme4* package (Bates et al., 2015). Data for saccadic precision, and saccadic reaction time were skewed and were both logarithmically transformed prior to analyses (best fitting transformation was selected for the data using Box-Cox Power transform). Moreover, extreme reaction times (> |2.5 SD|) were filtered prior to the analyses (2.86% of the trials were filtered). Visibility of the stimulus was fitted into the models as a centered continuous variable and, in Experiment 1, trial type (ipsilateral vs. contralateral TMS) was fitted into all the models as a repeated contrast coded (see Schad et al., 2020, for more detail) fixed factor. In the models, random effects were maximal (Barr et al., 2013). The exact degrees of freedom are difficult to determine for the *t* statistics estimated by mixed-effects models, leading to problems in determining *p*-values (Baayen et al., 2008). Statistical significance at the .05 level is indicated by the values of *t >* |1.96|. The data and the R-scripts are made available using the Open Science Framework https://osf.io/6wncf/

## RESULTS

### Experiment 1

#### General Overview of Task Performance

When a stimulus was presented, the participants reported seeing it (i.e., visibility rating > 0) in 87% of trials, on average. When no stimulus was presented, the participants reported seeing it in about 12% of the trials. TMS interfered with the participants’ conscious perception of the stimulus. When the visual stimulus was contralateral to TMS, the average (non-binarized) visibility rating was lower (*M* = 1.15, *SD* = 0.35) in comparison to ipsilateral trials (*M* = 1.48, *SD* = 0.35), *t*_11_ =−2.30, *p* = .042, *d* = 0.94 (see Figure 2B). The number of unconscious trials per participant was relatively high (*M* = 74, range = 9–145), indicating that, in general, consciousness was robustly suppressed by the TMS.

The proportion of saccadic reactions (to either stimulus location) is presented in Table 1 and illustrated in Figure 2A. The participants made a saccadic reaction, on average, in 69% of the trials in which a stimulus was presented, and they reported seeing it. The proportion of saccadic reactions was about the same between contralateral and ipsilateral TMS conditions: *t*_11_ =−0.76, *p* = .465, *d* = 0.11 (see Table 1 and Figure 2C). Of the saccadic reactions, 98% were made toward the correct location (i.e., the location in which the stimulus was presented) when the participants reported seeing the stimulus. The difference in the accuracy of saccadic reactions between contralateral and ipsilateral conditions was not statistically significant, *t*_11_ =−2.10, *p* = .060, *d* = 0.13 (see Table 1 and Figure 2D).

#### Ability to Make Saccades Towards Unseen Targets

When the participants reported not seeing the target (i.e., the lowest visibility rating), they made saccadic reactions to either target location in 7.7% of the trials in which a stimulus was presented. Given that the number of unconscious trials was high, this shows that, by and large, the initiation of saccades required conscious perception. However, we asked whether the participants made more saccadic reactions when a stimulus was presented (but they reported not seeing it), compared to catch trials (in which no stimulus was presented), indicating that the unconscious target influenced their behavior. Our aim was to examine the frequency of saccades to unconscious target stimuli separately during ipsilateral and contralateral TMS. However, because the participants made so few responses to targets, they reported not seeing, we combined the contralateral and ipsilateral conditions to maximize the number of unconscious trials. Three participants had only 0–2 saccadic reactions to unconscious target stimuli (total number of unconscious trials in which the target was presented in these participants were 9, 67, and 99); thus, they were excluded from the analyses (leaving a total of 718 unconscious trials in which the target was presented).

When the participants reported not seeing the target, they made saccadic reactions more often when a stimulus was present than when no target was presented, *t*_8_ =−2.43, *p* = .041, *d* = 0.97 (see Figure 3A). Moreover, when the participants made saccadic reactions, and reported not seeing the stimulus, they were able to shift their gaze toward the correct target location more often than predicted by chance (one-sample t test against 50%: *M* = 0.68, *SD* = 0.17), *t*_8_ = 3.31, *p* = .011, *d* = 1.73 (Figure 3B). Participants who were better at initiating saccadic reactions to unseen targets were also better at initiating saccades to the correct location, *r* = .83, 95% CI [.38, .96] (see Figure 3C).

We repeated the analysis of saccadic reactions to unconscious targets by calculating signal detection theoretical sensitivity (d’) and criterion (c). Although sensitivity to reacting to the unconscious target was low (*M* = 0.50, *SD* = 0.51), participants were able to discriminate the stimulus from the catch trials in which they reported not seeing any target (one-sample t-test against 0: *t*_8_ = 2.96, *p* = .018, *d* = 0.99). The response criterion was relatively high (M = 1.49, *SD* = 0.35), indicating that the participants were conservative in making saccadic reactions to targets they reported not seeing (i.e., the participants attempted to minimize false alarms).

To further elucidate the relationship between conscious perception and saccadic reactions, we calculated the participants’ sensitivity (d’) and criterion (c) for *reporting the conscious perception* of the target. The participants’ sensitivity to reporting conscious perception of the target (i.e., d’: *M* = 2.54, *SD* = 0.70) showed a moderate correlation with the proportion of correct saccadic answers when participants reported not seeing the target, *r* = .55 95% CI [-.29, .89]. This indicates that the participants who were less sensitive to subjectively detecting the target were less successful at launching saccades toward the target they reported not seeing (Figure 4A). Participants’ criterion for reporting conscious detection of the target (i.e., c: *M* = 0.40, *SD* = 0.28) also showed moderate correlation with the proportion of reactions toward the correct direction, *r* = .47, 95% CI [-.29, .86] (see Figure 4B). This indicates that a higher response criterion was associated with a better ability to unconsciously launch saccades toward the correct target location when a saccadic reaction was made. As the proportion of saccadic reactions (in either direction) correlated strongly with the proportion of correct saccades (Figure 3C), we only reported correlations with correct answers.

The sensitivity of consciously perceiving the stimulus showed a moderate correlation with the sensitivity of making saccadic reactions when the target was presented but reported as unseen, *r* = .62, 95% CI [-.07, .91] (see Figure 4C for illustration). The criterion for reporting that a target was seen versus unseen showed a moderate correlation with the criterion for making saccadic reactions (*M* = 1.49, *SD* = 0.35), *r* = .46, 95% CI [-.30, .86] (see Figure 4D for illustration).

#### Precision of Saccadic Reactions and Saccadic Reaction Times

Descriptive statistics for the precision and reaction time of correct saccadic reactions are presented in Table 2. Here, precision refers to the distance from the saccade landing position to the target position. The linear mixed-effects model on the *precision of the saccade* (when the stimulus was presented and the reaction was correct; see Table 3) revealed a main effect of *visibility*, indicating that the distance between saccade landing location and stimulus location decreased as the visibility rating increased (see Figure 5A). The model did not show an effect of trial type (contralateral vs. ipsilateral) or the interaction between visibility and trial type.

The model on *saccadic reaction times* (when the stimulus was presented and the reaction was correct; see Table 4), showed that higher stimulus visibility was associated with faster saccadic reactions (see Figure 5B). The model did not reveal an effect of trial type (Contralateral vs. Ipsilateral), or interaction between visibility and TMS location.

### Experiment 2

The main challenge in interpreting the results of Experiment 1 was the small number of saccadic reactions to unconscious targets. Although this indicates that, in general, the saccadic eye movements were based on conscious vision, it also suggests that inferences about saccades to unconscious targets may be unreliable (as they are based on a small number of data). Moreover, it is possible that (not just reduced visibility of the target but also) V1 stimulation interfered with the participants’ ability to initiate saccades toward the targets. To further test the ability to initiate saccades toward targets individuals do not see, we ran an experiment in which stimulus visibility was manipulated using a metacontrast mask instead of TMS. Metacontrast masking is assumed to interfere with target-related feedback activity, while leaving feedforward activity (in V1, and other areas) largely intact (Breitmeyer and Ogmen, 2010; Macknik and Livingstone, 1998; Railo and Koivisto, 2012). If participants are able to initiate saccades toward targets they report not consciously seeing, we should observe reflexive saccades toward the reportedly unseen targets.

#### General Overview of Task Performance

The proportion of saccadic reactions (i.e., first saccade made after stimulus onset) to either target location is presented in Table 5, and the end location of the saccadic reactions is illustrated in Figure 6A. When a target was presented, the participants reported seeing it (visibility rating > 0), on average, in 80% of trials. This is very similar to Experiment 1. When no stimulus was presented, the participants reported seeing a target stimulus, on average, in 39% of the trials. The number of false alarms was clearly higher than in Experiment 1 and likely due to the fact that the visual mask made it difficult for participants to accurately detect the target.

#### Ability to Make Saccades Towards Unseen Targets

When no target was presented, participants made a saccadic response to either potential target location, on average, in 85% of trials in which they reported seeing a target, and in 66% of trials in which they reported not seeing it. This indicates that the salient visual masks attracted the gaze. From the perspective of unconscious processing, the key question is whether the participants launched saccadic eye movements more often when a target they reported not seeing was presented on the screen when compared to the catch trials. The analysis showed that there were no differences between reactions toward catch versus target trials, *t*_12_ =−0.61, *p* = .552, *d* = 0.10 (see Table 5/Figure 6B). However, when the target was presented but the participant reported not seeing it, they were nevertheless able to initiate a saccade toward the correct target location in higher than guessing-level accuracy (*M* = 0.63, *SD* = .20), *t*_12_ = 2.40, *p* = .033, *d* = 0.67 (see Figure 6C). Similar to Experiment 1, the proportion of saccadic reactions (to either stimulus location) when the participants reported not seeing a target and proportion of reactions to correct location had a high positive correlation with each other, *r* = .77, 95% CI [.38, .93] (see Figure 6D).

As in Experiment 1, sensitivity (*d*’) and criterion (c) were calculated from saccadic reactions (i.e., the participant made a saccade to either target location when the target was presented) when the target was reported unseen. We analyzed whether participants were able to unconsciously distinguish and react to the target by calculating whether the d’ differs from zero. The results showed that the participants were not statistically significantly able to discriminate unconscious targets from catch trials (*M* = 0.37, *SD* = 0.79), *t*_12_ = 1.71, *p* = .113, *d* = 0.47. The participants’ criterion for making a saccadic response were more liberal when compared to Experiment 1 (*M* =−0.34, *SD* = 0.79).

We calculated the participants’ sensitivity and criterion for *reporting conscious perception* of the target and examined whether they predicted saccadic reactions to unconscious targets across participants. Overall, sensitivity to *report conscious perception* of the target was high (*M* = 1.30, *SD* = 0.76), and the response criterion was relatively low (*M*=−0.21, *SD* = 0.60). As the reactions in either direction correlated strongly with reactions to correct direction each, only correlations with correct answers were analyzed. The participants’ sensitivity to reporting conscious perception of the target showed a weak correlation with the proportion of correct saccadic answers when participants reported not seeing the target, *r* = .35, 95% CI [-.52, .57]. The criterion for reporting conscious detection of the target showed no correlation with the proportion of correct saccadic answers, *r* = .03, 95% CI [-.53, .57]. The sensitivity of making saccadic reactions when the target was presented but reported unseen also showed no correlation with the sensitivity of consciously perceiving the stimulus, *r* = .05, 95% CI [-.51, .59]. Lastly, response bias (i.e., *c*) regarding whether stimulus was seen versus unseen showed a weak correlation with the response bias measure of making saccadic reactions, *r* = .15, 95% CI [-.43, .65].

#### Precision Saccadic Reactions and Saccadic Reaction Times

Descriptive statistics for the precision and reaction time of the saccadic reactions are presented in Table 6. The model on *saccade accuracy toward target location* (see Table 7) showed that the higher the visibility rating, the closer the saccades were to the target location (Figures 7A). Moreover, the model on *saccadic reaction times toward the target location* (see Table 8) showed that saccadic reaction times became faster as visibility rating increased (see Figure 7B).

**Figure 7.**
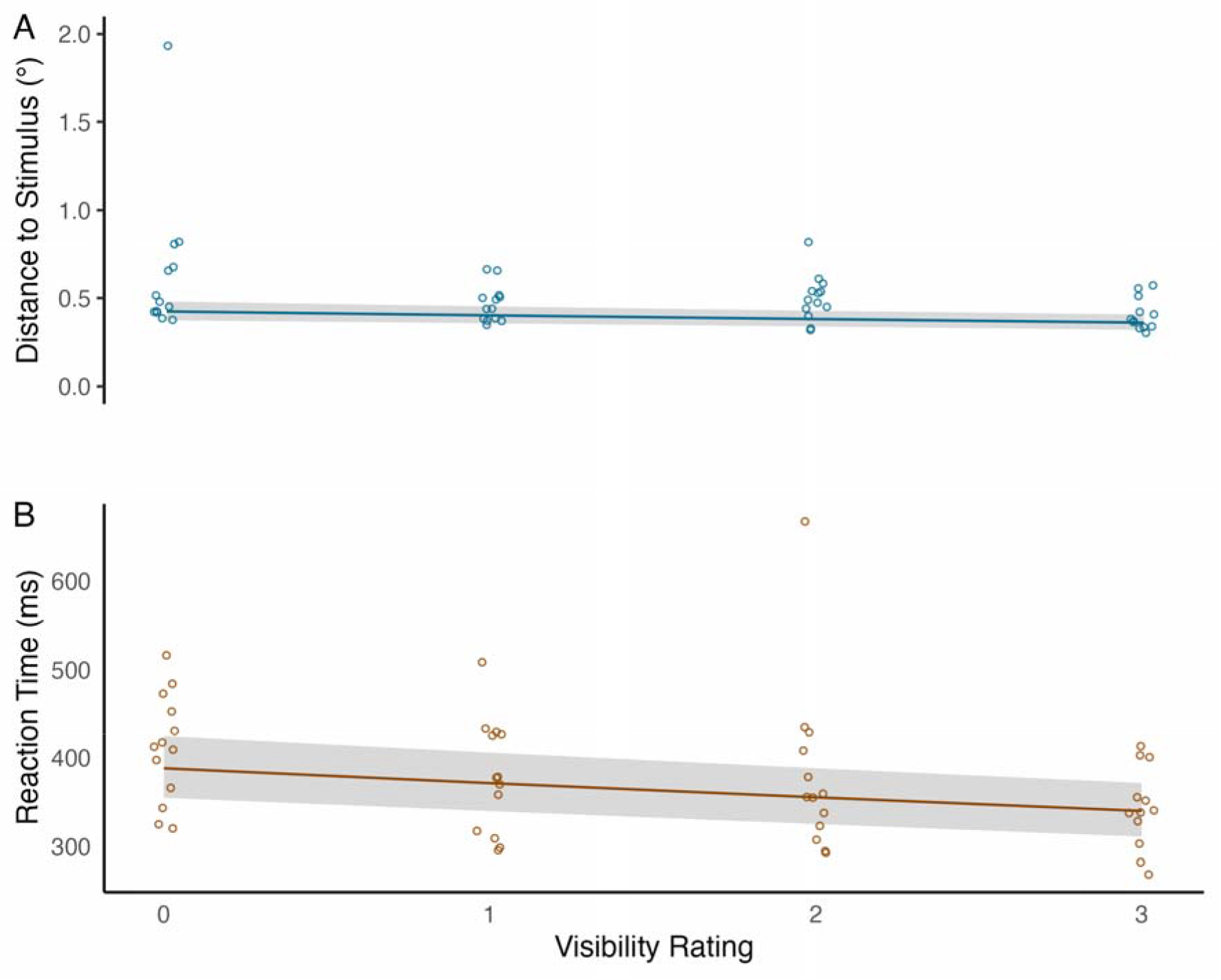
(A) Model estimates for the effect of visibility on saccade accuracy. (B) Model estimates for saccadic reaction times. For illustration purposes, the model estimates were back-transformed from log-values in both panels. Shaded area represents 95% CI. Circles represent the observed means for each participant.

**Table 6.** Descriptive statistics of the saccade accuracy and saccadic reaction times

| Visibility | Distance from stimulus location |  | RT (ms) |  |
| --- | --- | --- | --- | --- |
|  | *M* | *SD* | *M* | *SD* |
| Unseen | 0.52 | 0.99 | 403 | 108 |
| Seen | 0.44 | 0.33 | 353 | 93 |

*Note.* Distance from stimulus location is reported as visual degrees.

**Table 7.**
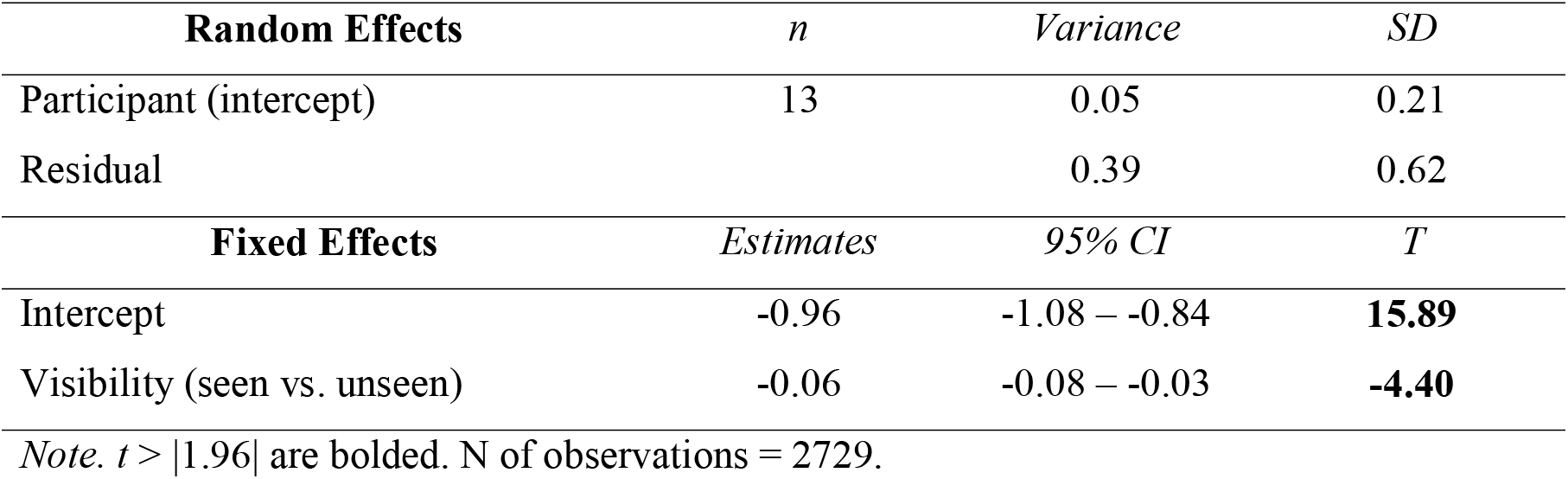
Model for distance of the end point of the saccadic answer from the stimulus location

**Table 8.** Model for saccadic reaction times

| **Random Effects** | *n* | *Variance* | *SD* |
| --- | --- | --- | --- |
| Participant (Intercept) | 13 | 0.03 | 0.16 |
| Residual |  | 0.04 | 0.21 |
| **Fixed Effects** | *Estimates* | *95% CI* | *t* |
| Intercept | 5.88 | 5.80 – 5.98 | **130.19** |
| Visibility | -0.05 | -0.06 – -0.04 | **-10.35** |

*Note.* $t > |1.96|$ are bolded. N of observations = 2673.

## DISCUSSION

In the present study, we tested neurologically healthy participants’ ability to orient their gaze toward visual targets that they reported not consciously seeing. The results were twofold. First, the *initiation* of saccades toward the target was largely based on conscious perception, and weaker conscious perception was associated with slower and less precise saccades. The results of the TMS experiment indicated that in a small number of trials, the participants were also able to initiate saccades toward targets they reported not seeing. The second main finding was that when the participants made saccadic reactions, and a target stimulus they were unaware of was presented, these saccades were made toward the correct target location. Experiment 2 using metacontrast masking largely replicated these findings, suggesting that the results cannot be attributed to disturbed processing in V1 in particular. In general, these results corroborate previous findings on blindsight patients (Barbur et al., 1988; Perenin and Jeannerod, 1978; Weiskrantz et al., 1974; Zihl, 1980) and monkeys (Isa and Yoshida, 2021; Kato et al., 2011; Yoshida et al., 2008), showing that gaze shifts can be influenced by unconscious information and may not depend on V1.

The present results replicate previous TMS-blindsight studies that have demonstrated that when the task is to simply detect the presence or localize a visual stimulus, blindsight-like behavior is observed (Christensen et al., 2008; Hurme et al., 2017; Railo and Koivisto, 2012; Ro, 2008, reviewed in Railo and Hurme, 2021). The present study is the first report that neurologically healthy participants can (on a small number of trials) initiate saccades toward target stimuli they report not seeing, and that this process may not depend on V1. The closest to the current study, Crouzet et al. (2014, 2017) studied whether saccades can be initiated toward a visual target that participants report not seeing. They observed that while participants could make accurate fast saccades to visual targets whose visibility was disturbed by visual masking, these responses accompanied a conscious perception of the stimulus. In the studies by Crouzet et al., the participants were asked to search for a target among distractor stimuli. The simpler task we employed in the present study may have been crucial to observing unconsciously guided saccades.

Although our results suggest that saccades can sometimes be initiated toward stimuli that individuals are unconscious of, it is very difficult to verify that participants indeed did not have any phenomenological experience related to the presented stimulus, even though they report not seeing it (Campion et al., 1983; Phillips, 2021). We used a graded visibility rating scale and defined unconscious stimuli as the lowest rating on a four-step scale because earlier reports of unconscious, blindsight-like capacity may have been confounded by the use of dichotomous rating scales (Koivisto et al., 2021; Mazzi et al., 2016; Overgaard, 2011). Despite the use of a graded rating scale, our analysis suggests that in Experiment 1, the reported unconscious saccades were nevertheless based on weakly conscious perception; participants who showed higher sensitivity to *consciously* detect stimuli also showed the highest “reportedly unconscious” capacity. This suggests that stimuli that participants reported as “absolutely not seen” may be associated with a weak degraded visual experience. This is consistent with research on “unconscious” saccadic reactions in monkeys (Isa and Yoshida, 2021; Yoshida and Isa, 2015). Similar correlations with conscious perceptual sensitivity and alleged “unconscious” perception have also been reported previously (Koivisto et al., 2021; Railo et al., 2021). Interestingly, a similar pattern was not observed in Experiment 2. This could be because TMS to V1 induces suboptimal introspection (Peters et al., 2017), or because participants made more false-alarm responses in Experiment 2 compared to Experiment 1.

Given that we applied TMS pulses 100 ms after visual target onset, it is likely that the fastest “feedforward” stimulus-evoked responses in V1 were not affected by TMS. This makes direct comparison to blindsight patients difficult (as lesions affect both feedforward, and later feedback activity). Theories of (un)conscious vision suggest that the feedforward activity period may enable unconsciously guided behaviors (Dehaene et al., 2006; Lamme and Roelfsema, 2000; Vanrullen, 2007; VanRullen and Koch, 2003). Consistent with this, Hurme et al. (2017) showed that unconscious blindsight-like behavior was eliminated when TMS pulses were applied after a delay that coincided with the timing of feedforward activity in V1 (60 ms after stimulus onset). This suggests that, in the present study, the saccades guided by unconscious stimuli may have been triggered by the transient feedforward activity passing through V1. Similarly, metacontrast masking, employed in Experiment 2, is assumed to leave feedforward activity largely intact (Breitmeyer and Ogmen, 2010; Macknik and Livingstone, 1998; Railo and Koivisto, 2012).

In Experiment 1, we wanted to target V1 using TMS to conceptually replicate the blindsight observed in patients. The aim was to compare conditions in which TMS was applied to contralateral versus ipsilateral V1 locations (Hurme et al., 2019; Hurme et al., 2017; Railo and Koivisto, 2012). That is, the ipsilateral TMS condition was originally included as a control condition, as we aimed to stimulate the same area while varying the stimulus laterality. However, the results revealed surprisingly small differences between the ipsilateral and contralateral conditions. In the present study, the stimulated V1 location in the left and right hemispheres was determined based on functional retinotopic mapping, whereas in our previous studies, TMS was typically targeted based on anatomy (Hurme et al., 2017, 2019; Railo and Koivisto, 2012). Possibly due to this difference, the V1 representation of the stimulus location was close to the interhemispheric fissure in the majority of the participants, meaning that stimulation could not be targeted specifically to either the contralateral or ipsilateral hemispheres. In other words, it is likely that the contralateral V1 was also stimulated in the ipsilateral condition. Furthermore, a notable methodological difference is that in our previous studies, we were able to use stronger visual stimuli than in the present study (e.g., Hurme et al., 2017, 2019), which might have enabled more selective suppression of contralateral stimuli.

Due to the small difference in performance in the contralateral and ipsilateral conditions, the evidence for the role of (retinotopically defined) V1 was not as strong as expected. Moreover, although retinotopic mapping ensured that we likely targeted V1, it is possible that V2 and V3 were also stimulated because these areas are often close to V1 (Salminen-Vaparanta et al., 2012; Thielscher et al., 2010). However, based on the data of the present experiment, we cannot tell whether the magnitude of the stimulation in these areas was strong enough to cause any behavioral effects, a limitation that is true for all TMS experiments targeting V1 (e.g., Salminen-Vaparanta et al., 2012). Further, in experiments conducted with patient and nonhuman primates, the lesions rarely limit specifically to V1. Thus, in this respect, the results remain conceptually compatible with previous studies on patients.

The TMS experiment rests on the assumption that we interfered with visual stimulus-related processing in V1. However, it could also be argued that saccadic reactions were influenced by TMS because the pulses disturbed processing in the fovea: possibly, the TMS pulses disturbed the *disengagement* of gaze from the central fixation cross. Although this counterargument cannot be completely refuted, several factors stand against this conclusion. First, compared to parafoveal targets, foveal representation is more difficult to disturb with TMS (e.g., because the representation of the fovea is larger, and thus more difficult to inhibit). Second, and most importantly, the results unequivocally showed that when participants consciously perceived the target stimulus, they were also able to make accurate saccades toward it (Figure 3B). This implies that the decrease in saccadic performance was due to suppressed visibility (not just the difficulty of disengaging from fixation). Third, the fact that similar effects were observed in the metacontrast masking experiment suggests that the results reflect reduced visibility (metacontrast masks do not affect processing in the fovea).

We used small visual stimuli (about 0.20 visual degrees) in the present study to maximize the chances of complete visual suppression by TMS (Experiment 1: *M* = 74 suppressed trials/participant, range = 9–145; Experiment 2: *M* = 61 suppressed trials/participant, range = 13–114). Despite the reasonably high number of unconscious trials, the number of saccadic reactions to unconscious targets was low (when compared to the false-alarm rate). This suggests that the initiation of saccades was largely based on conscious perception. In Experiment 1, two participants had to be excluded from the analyses because they were not able to make saccades towards unseen stimuli despite a high number of unconscious trials (*N*_unseen_ = 99 and 67). There are at least two possible explanations for why this happened. First, it is possible that these participants were cautious of making saccades towards unseen targets (i.e., their response criterion was very high) and, consequently, failed to perform in the task as intended. Second, it is possible that TMS suppression was higher with these participants than with other participants, and thus interfered more severely with their ability to make saccades.

It is possible that larger, or more meaningful (e.g., emotional) visual stimuli could have enabled a better ability to initiate saccades toward unconscious stimuli. Although saccadic reaction times were relatively fast in the present study (300–400 ms), they were not *reflexively* fast: express saccades can have onset latencies as fast as 100 ms (Kingstone and Klein, 1993; Marino et al., 2015; Schiller et al., 2004), and it is likely that the stimuli used in the present study were too small to trigger express saccades. Furthermore, in our study, decreased conscious visibility was associated with slower saccadic reactions, suggesting that saccades toward targets whose visibility was suppressed were not based on reflex-like circuits.

It is also worth noting that in many previous studies in patients (e.g., Zihl, 1980; Savina and Guitton, 2018) or nonhuman primates (Mohler and Wurtz, 1977; Yoshida et al., 2008), the participants performed clearly larger saccades than in the present study. This makes comparison of studies difficult. We used 2° eccentricity targets because in our experience, early visual cortical representation with higher eccentricity is more difficult to reach using TMS. Studies in patients and nonhuman primates also enable the use of larger stimuli, which likely increases the chances of observing saccades that are initiated based on unconscious information.

In sum, our results show that neurologically healthy humans can sometimes orient their gaze toward stimuli they subsequently report not consciously perceiving. However, such actions are less precise, slower, and clearly less frequent when compared to stimuli that are consciously perceived. Our results suggest that the ability to make saccadic eye movements toward unconscious stimuli may be at least partly independent of V1. The present results are important because previous reports of the ability to launch saccades toward unconscious stimuli have been based on patients and nonhuman primates with permanent V1 lesions. That similar capacity may be present even in neurologically healthy individuals suggests that it is not due to neural reorganization that often accompanies lesions but a property of the neurologically intact human brain.

## Acknowledgements

This research was supported by a grant from the Ella and Georg Ehrnrooth Foundation awarded to Henri Olkoniemi, and a grant from the Academy of Finland (grant #308533) awarded to Henry Railo. We would like to thank Teemu Laine for technical help with Experiment 1.

## Notes

https://osf.io/6wncf/

### Competing Interest Statement

The authors have declared no competing interest.

### Summary of Updates

Minor layout changes, black and white figures are changed to color figures.

